# Cystinosin is involved in Na^+^/H^+^ Exchanger 3 trafficking in the proximal tubular cells: new insights in the renal Fanconi syndrome in cystinosis

**DOI:** 10.1101/2025.02.12.637793

**Authors:** Veenita Khare, Jean-Claude Farre, Celine Rocca, Mouad Ait Kbaich, Cynthia Tang, Xuan Ma, Kavya Beiderman, Ioli Mathur, Rafael A. Badell-Grau, Anusha Sivakumar, Rola Chen, Sergio D. Catz, Stephanie Cherqui

**Affiliations:** Department of Pediatrics, University of California, San Diego, La Jolla, California, USA; Division of Biological Sciences, University of California, San Diego, La Jolla, California, USA; Department of Molecular Medicine and Molecular and Cellular Biology, The Scripps Research Institute, La Jolla, California, USA

**Keywords:** cystinosis, renal Fanconi syndrome, cystinosin, Na^+^/H^+^ Exchanger 3, proximal tubular cells, hematopoietic stem cells

## Abstract

Cystinosis is a systemic lysosomal storage disease resulting from a defective *CTNS* gene, leading to the accumulation of cystine in all organs. Despite the ubiquitous expression of cystinosin, the renal Fanconi syndrome (FS) is the first manifestation of cystinosis that presents early in life of the patients while other complications appear years later. Additionally, the cystine reduction therapy, cysteamine, does not prevent the FS. While the matter is still unresolved, it is apparent that specific function(s) of cystinosin in the proximal tubular cells (PTCs) beyond cystine transport explain the early tubular defects in cystinosis. Here, we report a novel interaction of cystinosin with the sodium/hydrogen (Na^+^/H^+^) exchanger proteins in the endosomes in both yeast and mammalian cells. One isoform of Na^+^/H^+^ exchanger, NHE3, is a major absorptive sodium transporter at the apical membrane of the proximal tubules. Cystinosin was found to play a significant role in NHE3 subcellular localization, trafficking, and resulting sodium uptake in PTCs. Interestingly, introduction of *CTNS* successfully rescued these defects in *CTNS*-deficient PTCs, whereas *CTNS-LKG*, the lysosomal and plasma membrane isoform of cystinosin, did not. NHE3 mislocalization was confirmed in *Ctns^-/-^* mice and cystinosis patient kidney. Interestingly, transplantation of wild-type hematopoietic stem and progenitor cells in *Ctns^-/-^* mice restored NHE3 expression at the brush border membrane. This study uncovers a new role of cystinosin in the trafficking of NHE3 in the PTCs that is evolutionary conserved, offering new insights in the pathogenesis of the renal FS in cystinosis and potential new therapeutic avenue for this pathology.

**Significance:** This study reveals a new role of cystinosin in the trafficking of NHE3, the Na^+^/H^+^ exchanger in the proximal tubular cells (PTCs) in the kidney. NHE3 contributes to the majority of renal sodium absorption and is essential for maintaining the homeostasis of the PTCs. Thus, this study offers new molecular insights in the endosomal trafficking of NHE3 and provides a new mechanism that may explain the underlying pathogenesis and early onset of the renal Fanconi syndrome in cystinosis. Additionally, these findings provide potential new drug targets for this pathology and are also relevant to our ongoing stem cell gene therapy clinical trial for cystinosis, offering valuable information on the potential impact of hematopoietic stem cell transplantation on renal Fanconi syndrome.

## Introduction

Cystinosis is an autosomal recessive lysosomal storage disorder caused by mutations in the *CTNS* gene which encodes for the cystine transporter protein, cystinosin (1). While *CTNS* is ubiquitously expressed and cystine accumulates in all tissues, the kidney is the primary organ affected which shows the earliest manifestation of the disease phenotype with renal Fanconi syndrome (FS) at 6-18 months of age (2). The renal FS manifests by generalized dysfunction of the proximal tubules resulting in the ineffective reabsorption of essential nutrients such as glucose, uric acid, phosphate, amino acids, and bicarbonate (HCO3^-^) (2, 3). Interestingly, the onset of FS in cystinosis patients occurs almost concurrently with dedifferentiation of proximal tubular cells (PTCs) often prior structural abnormalities, such as cystine crystal formation or accumulation can be detected, suggesting that the pathophysiology of cystinosis involves mechanism beyond cystine accumulation (4). Cystinosis is currently the leading cause of inherited renal Fanconi syndrome (FS) in children, representing up to 20% of patients with hereditary tubular disorders (2, 3). Patients affected with cystinosis also develop a progressive loss of glomerular function eventually leading to renal failure in the second or third decade. The non-renal manifestations of cystinosis have become apparent, and they develop severe impairment of other organs including the heart, thyroid, muscle, pancreas, eye, and central nervous system (5).

The mouse model of cystinosis, the *Ctns^-/-^* mice, accumulates cystine and cystine crystals, pathognomonic of cystinosis, in all tissues (6). The pure strain of C57BL/6 *Ctns^-/-^* mice develop kidney pathology, in particular, a renal Fanconi syndrome, albeit less severe than in humans, and eventually end-stage renal failure (7). We previously reported that syngeneic wild-type (WT) hematopoietic stem and progenitor cell (HSPC) transplantation in myeloablated *Ctns^-/-^* mice resulted in HSPC-derived engraftment within the kidney, improvement of the renal Fanconi syndrome, and long-term preservation of kidney function and structure (8, 9). The mechanism underlying this therapeutic effect involves the differentiation of WT HSPCs into macrophages, which provide “healthy lysosomes” carrying the functional protein cystinosin to the host diseased cells via extension of tunnelling nanotubes (TNTs) that can even cross the basement membrane to deliver cystinosin to the PTCs (10). We developed an autologous HSPC transplantation after ex vivo gene-correction of the HSPCs using a lentiviral vector containing the *CTNS* cDNA (11, 12), and conducted a phase 1/2 clinical trial including six adult patients (ClinicalTrials.gov Identifier: NCT03897361) (13). While the impact of the therapy on the kidney disease could not be evaluated in this trial because the patients had too advance kidney disease or kidney transplants, the overall outcomes are promising.

The current substrate reduction therapy for cystinosis, cysteamine, delays disease progression but does not prevent renal Fanconi syndrome, eventually leading to end-stage renal failure (14). This strongly suggests that cystinosin has another specific function in the proximal tubular cells beyond lysosomal cystine-proton cotransporter. In recent years, studies have revealed novel roles of cystinosin in PTCs such as its involvement in autophagy (15–18), including chaperone-mediated autophagy (19), TFEB expression (20), and inflammation (21). Notably, none of these defective pathways were rescued by cysteamine (15, 19, 20). Furthermore, there are two isoforms of cystinosin protein due to alternate splicing of *CTNS* within exon 12 resulting in the exclusion of the GYDQL motif at the carboxyl-terminal and the longer *CTNS-LKG* isoform. This variant is expressed in various cellular compartments including lysosomes, plasma membrane, endoplasmic reticulum, and other cytoplasmic vesicles, but its specific function is unknown (22).

In this study, we report a novel interaction between cystinosin and members of the sodium-hydrogen (Na^+^/H^+^) exchanger family (NHEs). The Na^+^/H^+^ exchanger genes (SLC9 family), which include 10 isoforms with different tissue and cellular localization, play an essential role in maintaining cellular homeostasis, regulating factors such as cell pH, volume, and sodium concentration (23–25). The Na^+^/H^+^ exchanger isoform 3 (NHE3) is of particular interest because of its localization at the apical membrane of proximal tubules in the kidney and its role in the absorption of water and sodium (26). NHE3 is also crucial for preventing metabolic acidosis by enabling bicarbonate reabsorption (27). We showed that cystinosin and NHE3 interact and colocalize at the endosomes in the yeast model and human PTCs, and that cystinosin is necessary for its trafficking to the brush border in human PTCs. We also demonstrated that cystinosin, but not cystinosinLKG, is able to rescue the defective vesicular trafficking, and sodium and albumin uptake in *CTNS*-deficient PTCs. Notably, mislocalization of NHE3 was confirmed in *Ctns^-/-^* murine and cystinosis patient kidneys, and WT HSPC transplant was able to rescue its localization to the brush border. Together, this study provides a new role of cystinosin in the PTCs, supporting a molecular mechanism underlying the renal FS in cystinosis and supporting new therapeutic targets for this pathology.

## Results

### Ers1 colocalizes and interacts with Nhx1 in the endosomes in yeasts

Ers1 is well conserved and shares structural and functional similarities with human cystinosin (Fig. S1) (28). A high-throughput screen using modified split-ubiquitin technology in *S. cerevisiae* previously identified an interaction between the yeast cystinosin homolog, Ers1, and the NHE homolog protein, Nhx1 (29). The functions of mammalian NHE exhibit conservation in yeast as well, where Nhx1 transports both K⁺ and Na⁺, facilitating vacuolar Na⁺ sequestration by coupling its movement to the proton gradient generated by vacuolar H⁺-ATPase, thereby playing a role in pH regulation within luminal and cytoplasmic compartments and enhancing salt and osmotic tolerance (30–32). To confirm the interaction between Ers1 and Nhx1, we used the methylotrophic yeast *Pichia pastoris*, where the two cystinosin isoforms found in higher eukaryotes, exist but are encoded by two distinct genes: Ers1S (short), homologous to cystinosin, and Ers1L (long), homologous to cystinosinLKG, unlike *S. cerevisiae* that only has the short isoform, Ers1S (Fig. S1). Like *S. cerevisiae*, both forms of *P. pastoris* Ers1 structure are well conserved but lack the luminal N-terminal domain (NTD) (Fig. S1). This suggests that *P. pastoris* has evolved a divergent mechanism to achieve the same functional outcome, making it an excellent model to study cystinosis. To investigate the potential interaction, we performed the in vivo live-cell imaging assay, Bimolecular Fluorescence Complementation (BiFC), for direct visualization and localization of protein-protein interactions. Two non-fluorescent fragments of the Venus fluorescent protein (Venus-N and Venus-C) were fused with Ers1S and Nhx1 to create Ers1S- Venus-N (Ers1S-VN) and Nhx1-Venus-C (Nhx1-VC), respectively. Fluorescent signal generated by complementation of these two non-fluorescent fragments was observed in small vesicles in the cytoplasm and confirmed the interaction of Ers1S and Nhx1 in these cellular compartments (Fig. 1A). Budding yeast possess a minimal endomembrane system lacking a distinct early endosome (EE), and the trans-Golgi network (TGN) serves as the EE (33). The majority of Ers1S-VN+Nhx1-VC dots colocalized with the late endosome (LE) marker, Vps8, with some partial colocalization with the TGN/EE marker, Sec7, suggesting trafficking of Ers1S-VN+Nhx1-VC between the TGN/EE and LE (Fig. 1A). These results are consistent with the endosomal localization of Nhx1 reported in *S. cerevisiae* (34). We then examined the subcellular localization of Ers1S using Ers1S-GFP fusion protein in ΔErs1S *P. pastoris* strain and conducted pulse-chase experiments with the dye FM4-64, which enters the yeast via endocytosis and selectively stains endocytic intermediates, including early and late endosomes, as well as the vacuole, with red fluorescence (35). By fluorescence microscopy, we observed Ers1S-GFP colocalizing with endocytic intermediates labeled with FM4-64 after a short pulse-chase (45 minutes) but showed surprisingly no localization with the vacuole at either 45 or 105 minutes of the chase (Fig. 1B). We speculated that Ers1 might play a role in Nhx1 localization and/or function at the endosome, impacting proper trafficking within the endocytic pathway. Hence, we investigated whether Ers1 contributes to cellular trafficking of Nhx1 by generating various mutants of Ers1S/L, including the double *ers1* deletion strain (Δ*ers1S* Δ*ers1L*), and visualizing Nhx1-GFP and Vps8-2xmCherry, the LE marker. As expected, Nhx1 colocalized with Vps8 in wild-type cells; however, colocalization was also observed in single *ers1* (Δ*ers1S* or Δ*ers1L*) and double (Δ*ers1S* Δ*ers1L*) mutant cells. Interestingly, two major differences were observed for Nhx1 localizations in the Δ*ers1S* Δ*ers1L* double mutant, first Nhx1 protruded beyond Vps8 (yellow arrows; Fig. 1C) and second a large fraction of Nhx1 localized into the vacuolar matrix (yellow circle; Fig. 1C). This suggests that the double deletion of Ers1 (S and L) might affect late endosome/multivesicular body (MVB) morphology. In the absence of both Ers1 form, Nhx1 might be degraded by vacuolar proteases, consistent with Fig. 1C (yellow circle). With the goal of understanding the mechanism of Ers1’s trafficking role in Nhx1, we investigated the localization of Nhx1 in several single mutants implicated in vesicular trafficking in yeast, such as Δ*vps*1 (dynamin GTPase homolog implicated in several steps of the endocytic pathways, homolog of mammalian dynamin-1) (36), Δ*vps15* (member of phosphatidylinositol 3-kinase complex required for autophagy and endosomal vacuolar protein sorting, homolog of mammalian p150) (37), and Δ*ypt7* (GTPase required for vesicular fusion with the vacuole, homolog of mammalian RAB7) (38). No mislocalization of Nhx1 was observed in the absence of Ypt7 or Vps15. However, in cells lacking Vps1, Nhx1 accumulated around the vacuole/lysosome (yellow circle, Fig. 1D). This mislocalization in the *vps1* mutant is similar to what has been previously observed for Snc1 (mammalian homolog of vertebrate synaptic vesicle-associated membrane proteins (VAMPs) or synaptobrevins) (39), Vps10 (mammalian homolog of sortilin) (40), and Pep12 (mammalian homolog of syntaxins) (41) in yeast, confirming that Nhx1 traffics through the endosomal system. This suggests as well that Ers1 may function with Vps1 to recycle Nhx1 between the TGN/EE and late endosomes.

**Figure. 1.**
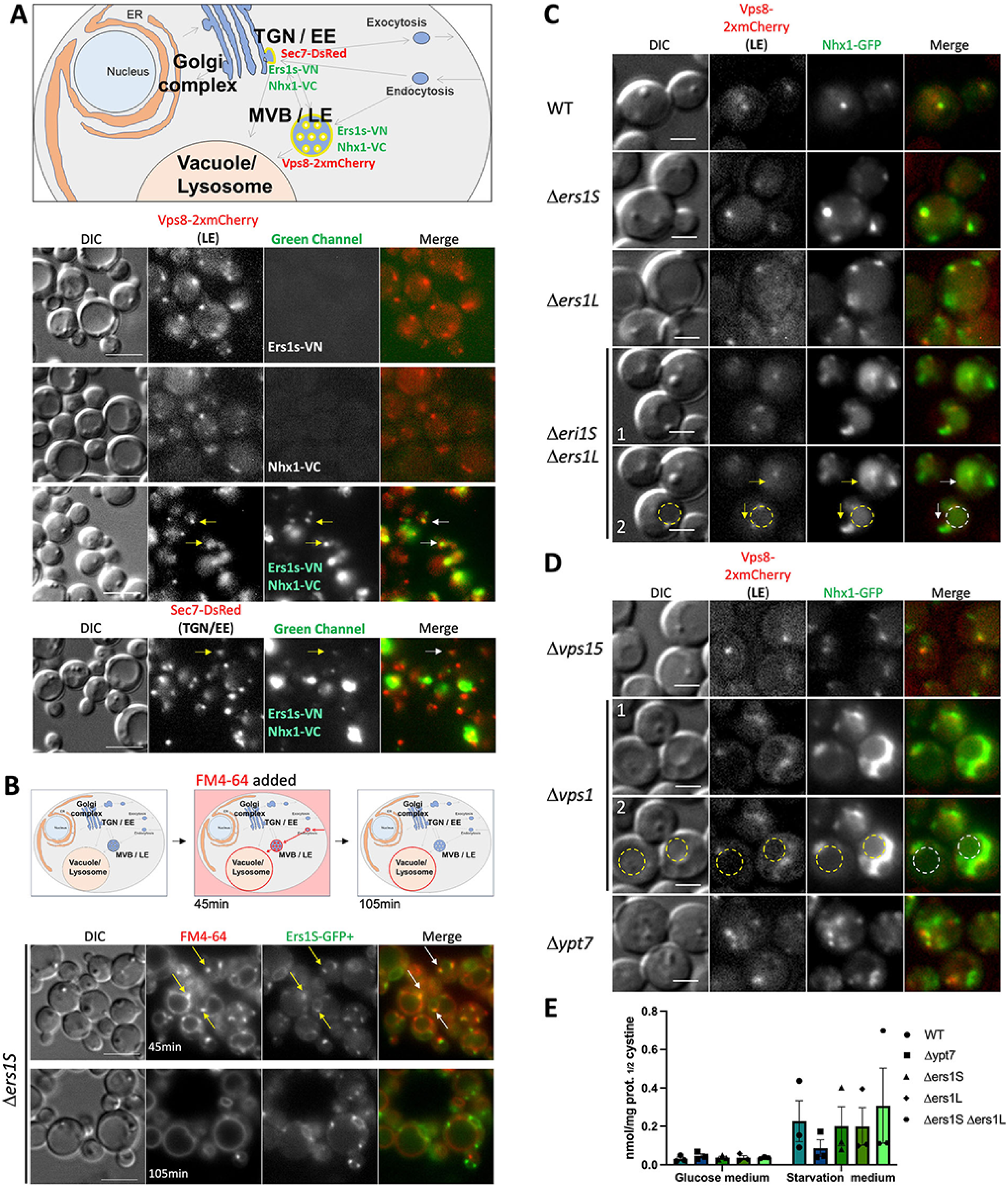
Ers1 colocalizes and interacts with Nhx1 in the endosomes in yeast cells. **(A)** Schematic representation of the BiFC assay in *P. pastoris* displaying an interaction between Ers1s and Nhx1, and their colocalization with the early and the late endosome marker, Sec7-DsRed (TGN/EE) and Vps8-2xmCherry (MVB/LE), respectively (**Upper panel**). BiFC assay performed shows that the interaction between Ers1s and Nhx1 occurs at the endosome, through colocalization with Vps8-2xmCherry and Sec7-DsRed (arrows). Negative controls, the single Venus moieties (Ers1s-VN and Nhx1-VC) do not fluoresce in the green channel, confirming their interaction (**Lower panel)**. **(B)** Schematic representation of the vital stain FM4-64 internalization and transport to the vacuole/lysosome in yeast. Pulse-chase of FM4-64 followed by a short incubation (45 min) stains endocytic intermediates, such as early and late endosome and the vacuole, whereas a longer incubation (105 min) labels mainly the vacuole (**Upper panel**). Fluorescent microscopy shows colocalization of Ers1S-GFP with endocytic intermediates labeled with FM4-64 (45min) but no colocalization with the vacuole at either 45 or 105min. Cells deleted for *ERS1S* gene were complemented with a plasmid comprising Ers1S-GFP under regulation of the Ers1 promoter. Arrows indicate colocalization of Ers1S-GFP with endocytic intermediates (**Lower panel)**. **(C & D)** Localization of Nhx1-GFP expressed under the Nhx1 promoter in different mutant strains (dashed circles indicate the vacuole and arrows indicate Nhx1-GFP localization protruding from the late endosome). Panels 1 and 2 show the same images, with and without annotations. Bars: 5 mm in (**A & B**), 2 mm in (**C & D**). **(E)** Cystine content was measured by mass spectrometry in WT, Δ*ypt7* yeast strains and in Ers1S and L knockout strains as well as in the double knockout mutants (both Ers1S and L) in both regular glucose culture medium and under starvation conditions. Δ*ypt7* strain was used as a negative control, as its absence results in low amino acid content during nitrogen starvation due to impaired autophagosome-vacuole fusion. Two-way analysis of variance (ANOVA) was used with comparison done between different cell lines under the same condition. The data in (E) represent the means and standard error of mean (SEM) from three biological replicates.

Finally, we aimed to confirm whether the deletion of Ers1 in *P. pastoris*, similar to *S. cerevisiae* (42), did not lead to cystine accumulation. We studied *P. pastoris* strains lacking Ers1S, Ers1L, or both, under both growth conditions (glucose medium) and nitrogen starvation conditions (starvation condition), where cells arrest the cell cycle and induce autophagy to recycle material for cell survival (Fig. 1E). Under growth conditions, we confirmed that *P. pastoris* behaved similarly to *S. cerevisiae*, with *ERS1* deletion strains not accumulating cystine. Surprisingly, the same result was observed under nitrogen starvation conditions. Nitrogen starvation in yeast induces autophagy and increases amino acid content in the vacuole. Altogether, these results support that Ers1 protein does not function as a cystine transporter in the yeast and is involved in the vesicular trafficking of Nhx1.

### Cystinosin interacts and localizes with NHE3 & NHE2, but not with NHE1

To investigate potential interaction between NHE3 and cystinosin in mammalian cells, we generated stable HEK293T cell lines with lentiviral vectors (LV) expressing *NHE3* fused with the green fluorescent protein (*GFP*; LV-NHE3-GFP), along with *CTNS* and *CTNS-LKG* fused with *DsRed* reporter gene (LV-CTNS-DsRed; LV-CTNS-LKG-DsRed). Subsequent immunoprecipitation (IP) assays using GFP or DsRed tags from whole cell lysates demonstrated distinct co-precipitation of NHE3-GFP with both cystinosin-DsRed and cystinosinLKG-DsRed, and *vice versa* (Fig. 2A). Additionally, positive colocalization patterns between cystinosin-DsRed and NHE3-GFP and between cystinosinLKG-DsRed and NHE3-GFP observed by fluorescent microscopy within distinct intracellular vesicles (Fig. 2B). Subsequently, we aimed to determine whether this interaction was specific to NHE3 so we examined two other isoforms, NHE1, expressed ubiquitously, and NHE2, expressed primarily in the gastrointestinal tract and kidney (25, 43, 44). Intriguingly, we found that NHE2-GFP, but not NHE1-GFP, co-precipitated with cystinosin-DsRed and cystinosinLKG-DsRed, and *vice versa* (Fig. 2C, Fig. S2). Altogether, these results demonstrated the interaction of cystinosin and NHE3 and NHE2 and underscored the specificity of cystinosin interaction with some Na^+^/H^+^ exchanger proteins.

**Figure. 2.**
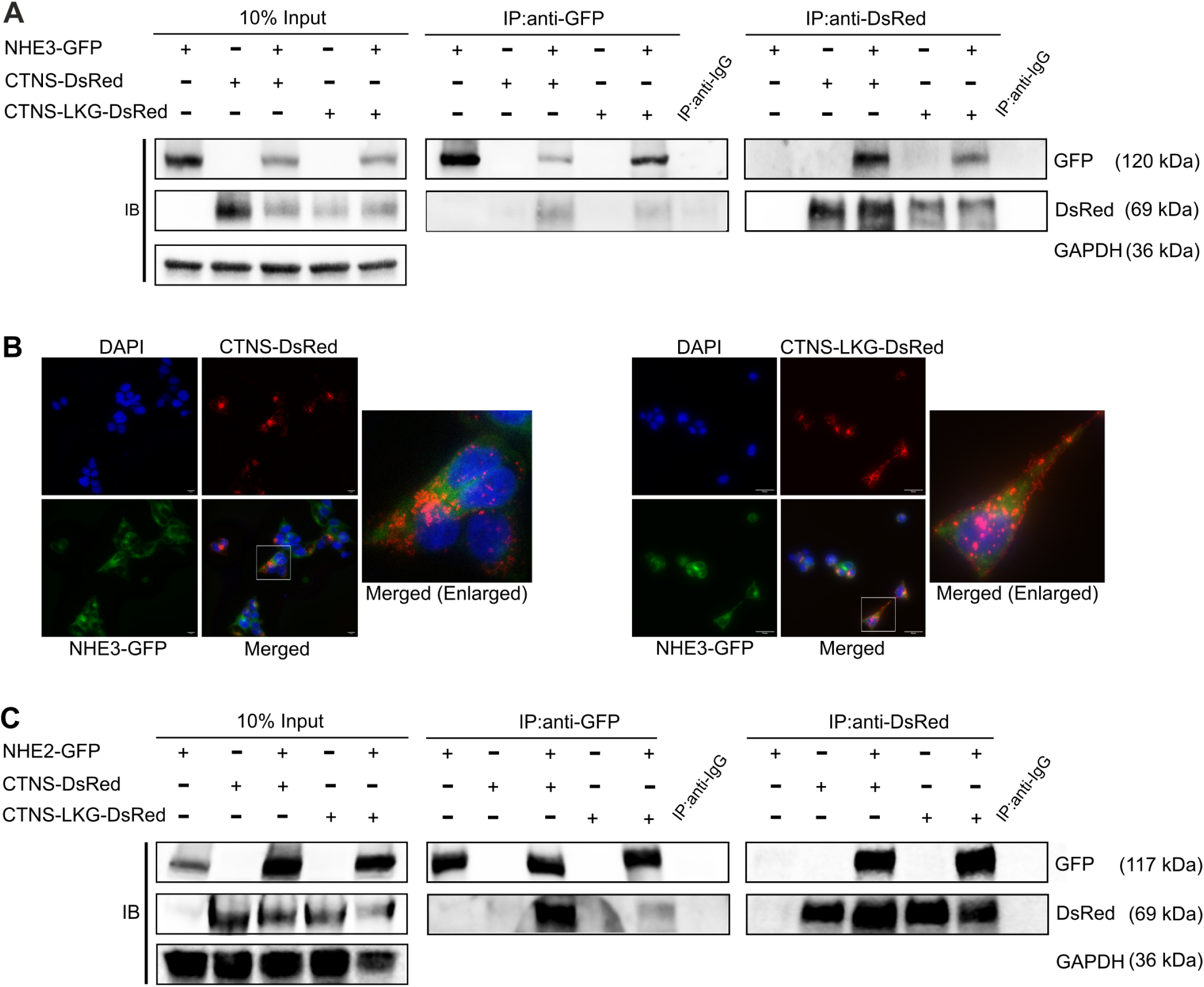
Cystinosin interacts with NHE3 and NHE2. (**A**) Immunoprecipitation assay performed in HEK293T cells stably expressing NHE3, cystinosin, and cystinosinLKG. Pull down using anti-GFP and anti-DsRed demonstrate the interaction between NHE3 and both cystinosin and cystinosinLKG. (**B**) Representative immunofluorescence images of HEK293T cells stably expressing NHE3-GFP and cystinosin-DsRed, and NHE3-GFP and cystinosinLKG-DsRed depicting colocalization of NHE3 with cystinosin, and NHE3 with cystinosinLKG in intracellular vesicles. Bars = 20 μm (left panel) and 50 μm (right panel) **(C**) Immunoprecipitation assay performed in HEK293T cells stably expressing NHE2, cystinosin, and cystinosinLKG showing interaction between NHE2 and both cystinosin and cystinosinLKG. For both (**A** & **C**) input lysates show proper expression of the proteins with glyceraldehyde-3-phosphate dehydrogenase (GAPDH) used as a loading control. Pull down with anti-IgG was used as a negative control. A representative image from three independent biological repeats is shown.

### *CTNS*^-/-^ PTC lines exhibit decreased expression, mislocalization of NHE3 and impaired sodium content and albumin uptake

To determine whether *CTNS* influences the expression of the NHE isoforms under investigation, we measured their endogenous expression levels in the proximal tubular cell line HK2 wild-type (WT) and *CTNS^-/-^* cells. At the mRNA level, quantitative real-time polymerase chain reaction (PCR) analysis did not reveal major changes in *NHE3* and *NHE1* transcripts in *CTNS*^-/-^ HK2 cell lines compared to WT (Fig. 3A). However, interestingly, significant increase in *NHE2* transcripts was observed in *CTNS*^-/-^ cell compared to WT. As expected, *CTNS* transcripts were not detected in *CTNS^-/-^* HK2 cells, which accumulate large amount of cystine (45), confirming their genetic status. Western blot analysis conducted on total cell extracts revealed a significant decrease in NHE3 expression as well as a significant increase in NHE2 expression in *CTNS*^-/-^ cells compared to WT cells. No alteration in the protein expression of NHE1 was observed (Fig. 3B).

**Figure. 3.**
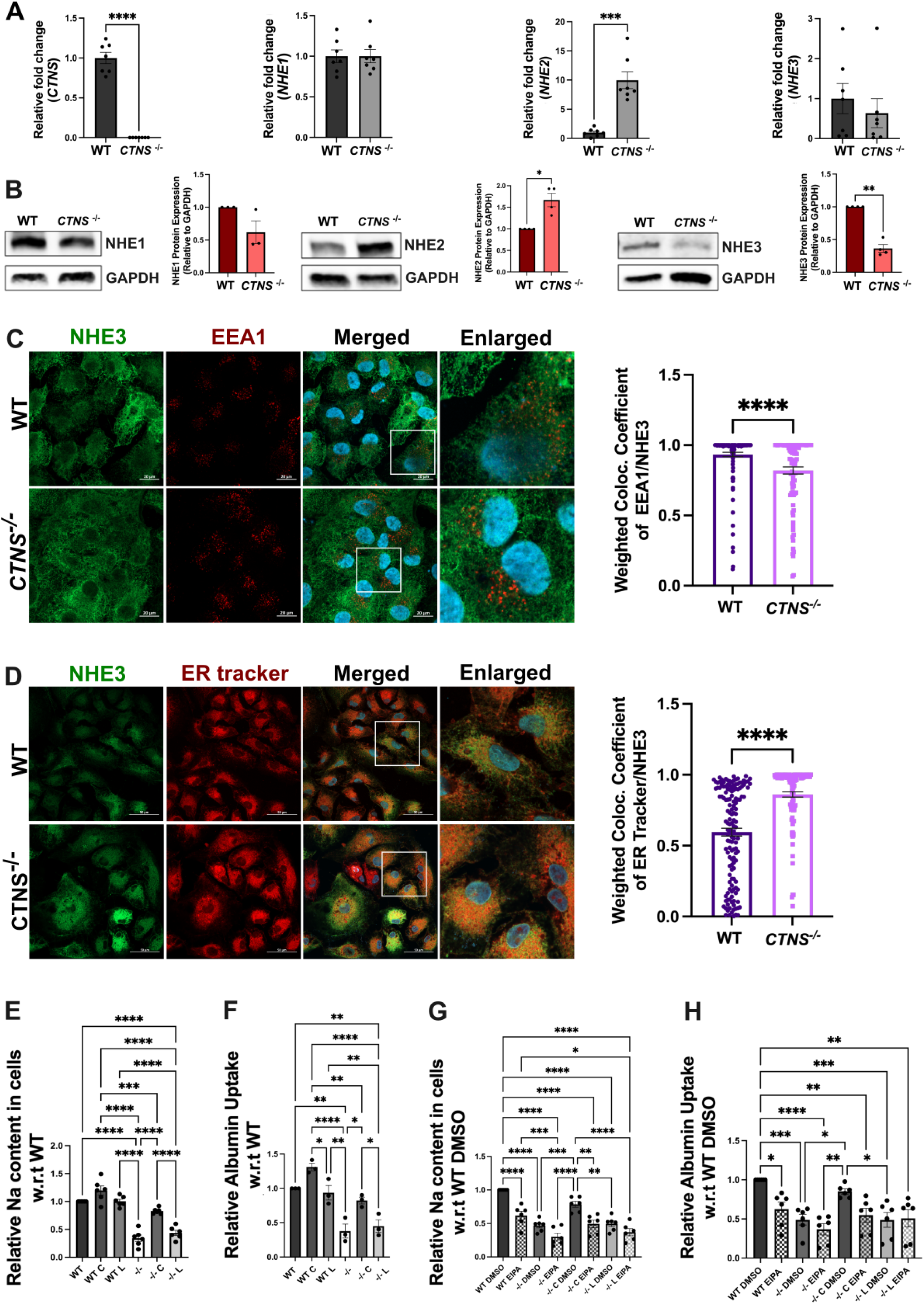
Cystinosin deficiency affects NHE3 expression, causing its mislocalization and impairing sodium and albumin uptake in human WT and *CTNS^-/-^* HK2 proximal tubular cells. (**A**) *CTNS*, *NHE1*, *NHE2*, and *NHE3* expression measured with quantitative polymerase chain reaction. (**B**) Representative Western blot analysis of NHE1, NHE2 and NHE3 expression with GAPDH used as a loading control. (**C, D**) Representative immunofluorescence images of colocalization of NHE3-GFP (green) with the early endosome marker EEA1(red) (**C**) and with the endoplasmic reticulum (ER tracker; red). (**D**) The corresponding quantification showed a defect in NHE3 trafficking that accumulated in the ER under *CTNS^-/-^* condition. Scale bar = 20 μm for both **C** and **D**. (**E and F**) Sodium (Na) and albumin uptake studies in WT and *CTNS^-/-^* HK2 cells (-/-), and in cells transduced with LV-CTNS (C) or LV-CTNS-LKG (L) measured using Sodium Green™ Tetraacetate and albumin555 by flow cytometry. (**G** and **H**) Na and albumin uptake in WT and *CTNS^-/-^* HK2 cells (-/-), and in *CTNS^-/-^* HK2 cells transduced with LV-CTNS (C) or LV-CTNS-LKG (L) treated with DMSO (vehicle), or the sodium transporter inhibitor (EIPA). For data in figure (A-B & E-H) each dot represents an independent biological replicate. The data in (C & D) represent the means and standard error of mean (SEM) from three biological replicates. Bar graphs are presented as the mean ± SEM and for (**A-D**) two-tailed Student’s t-test and for (**E**-**H)** One-way analysis of variance (ANOVA) was used, \**P* < 0.05; \*\**P* < 0.01; \*\*\**P* < 0.001; \*\*\*\**P* < 0.000.

To characterize the subcellular localization of NHE3 in PTCs, we transduced the WT and *CTNS^-/-^* HK-2 cells with LV-NHE3-GFP and studied its localization pattern with lysosomes (LAMP1), early endosomes (EEA1), Golgi (GM130), and endoplasmic reticulum (ER) (Fig. 3C-D, Fig. S3). We observed significant differences in the localization pattern of the NHE3 protein, being localized primarily in the ER and to a lesser extent in the lysosome, Golgi and early endosomes under *CTNS*^-/-^ condition as opposed to WT. These data suggest that the absence of cystinosin causes trafficking defects inducing the accumulation of NHE3 in the ER (Fig. 3C-D, Fig. S3).

Finally, we investigated the impact of the absence of cystinosin on NHE3 function. NHE3 is involved in sodium uptake in PTCs and plays a role in albumin endocytosis (46). To assess intracellular sodium content the cells were incubated with Sodium Green™ Tetraacetate for 8 mins. To induce the expression of megalin and cubilin in HK2 cells, we treated them with albumin555 for 16 hours as previously described (47, 48). Intracellular sodium and albumin uptake were then measured using flow cytometry analysis. We observed significantly lower intracellular sodium and albumin uptake in the *CTNS*^-/-^ cells compared to WT (Fig. 3E-F). Interestingly, the defect was rescued when the cells were transduced with LV-CTNS, but not with LV-CTNS-LKG. We also conducted these assays in the presence of the NHE3 inhibitor, EIPA (49, 50), which resulted in a significant decrease in sodium content and albumin uptake in WT HK2 cells. Notably, no significant decrease was observed in *CTNS^-/-^* HK2 cells treated or not with EIPA, and this effect persisted even after *CTNS* transduction in *CTNS^-/-^* HK2 cells treated with EIPA, thereby confirming the role of NHE3 (Fig. 3G-H). These results demonstrate the key role played by cystinosin in the regulation of NHE3 expression, localization and function.

### Cystinosin is involved in the cellular trafficking of NHE3 in PTCs

To confirm the defective trafficking of NHE3 in cystinosis PTCs, we studied the localization and expression of NHE3 protein in human healthy- (referred as Normal) and patient- (referred as CT) derived PTCs after transduction with LV-NHE3-GFP. NHE3 undergoes trafficking between the plasma membrane and endosomes (51). We hypothesized that the defective vesicular trafficking of Nhx1 observed in Ers1-defective yeasts might also occur in mammals. To further explore the vesicular dynamics of NHE3 and determine if its mislocalization resulted from defective trafficking, we measured its vesicular transport using pseudo-Total Internal Reflection Fluorescence microscopy (pTIRFM, oblique illumination). In pseudo-TIRFM, laser illumination is adjusted to impinge on the coverslip at an angle of incidence that is slightly less than the critical angle required for complete internal reflection. This technique facilitates imaging the cell surface by detecting the trafficking events in proximity to the plasma membrane while also imaging deeper areas in the cell than traditional TIRFM (the sample is illuminated up to ∼1 μm depth) (52). Thus, pTIRFM allows complete visualization of endosomes, lysosomes and other subcellular organelles but still maintaining the high signal-to-noise ratio of traditional TIRFM (52). In this experiment, vesicles were monitored for a period of 60 seconds. Representative images of normal, cystinotic patient derived PTCs and rescued cells are shown in Fig. 4A and associated movies 1-4. Dynamic studies show that cystinotic cells are characterized by a significant decrease in the number of NHE3^+^ organelles moving at high speed (>0.5 µm/sec) and a concomitant increase in the number of these organelles with no or restricted movement (<0.14µm/sec) in cystinotic cells (Fig. 4B-C). These data suggest defective NHE3 trafficking in patient-derived PTCs (Fig. 4A-B). Remarkably, transduction of CT PTCs with LV-CTNS but not with LV-CTNS-LKG significantly reduced the number of NHE3^+^ vesicles showing restricted movement in cystinotic cells (Fig. 4B), suggesting that cystinosin, but not cystinosinLKG, regulates NHE3 trafficking (Fig. 4A-C). Furthermore, sodium content in the CT cells was significantly lower in comparison to healthy cells, confirming a significant defect in sodium uptake in cystinosis patient PTCs compared to control (Fig. 4D). This defect was restored upon transduction with LV-CTNS, but not LV-CTNS-LKG. Altogether, these results show that cystinosin, but not cystinosinLKG, is involved in NHE3 vesicular trafficking and subcellular localization, impacting NHE3 function in PTCs.

**Figure. 4.**
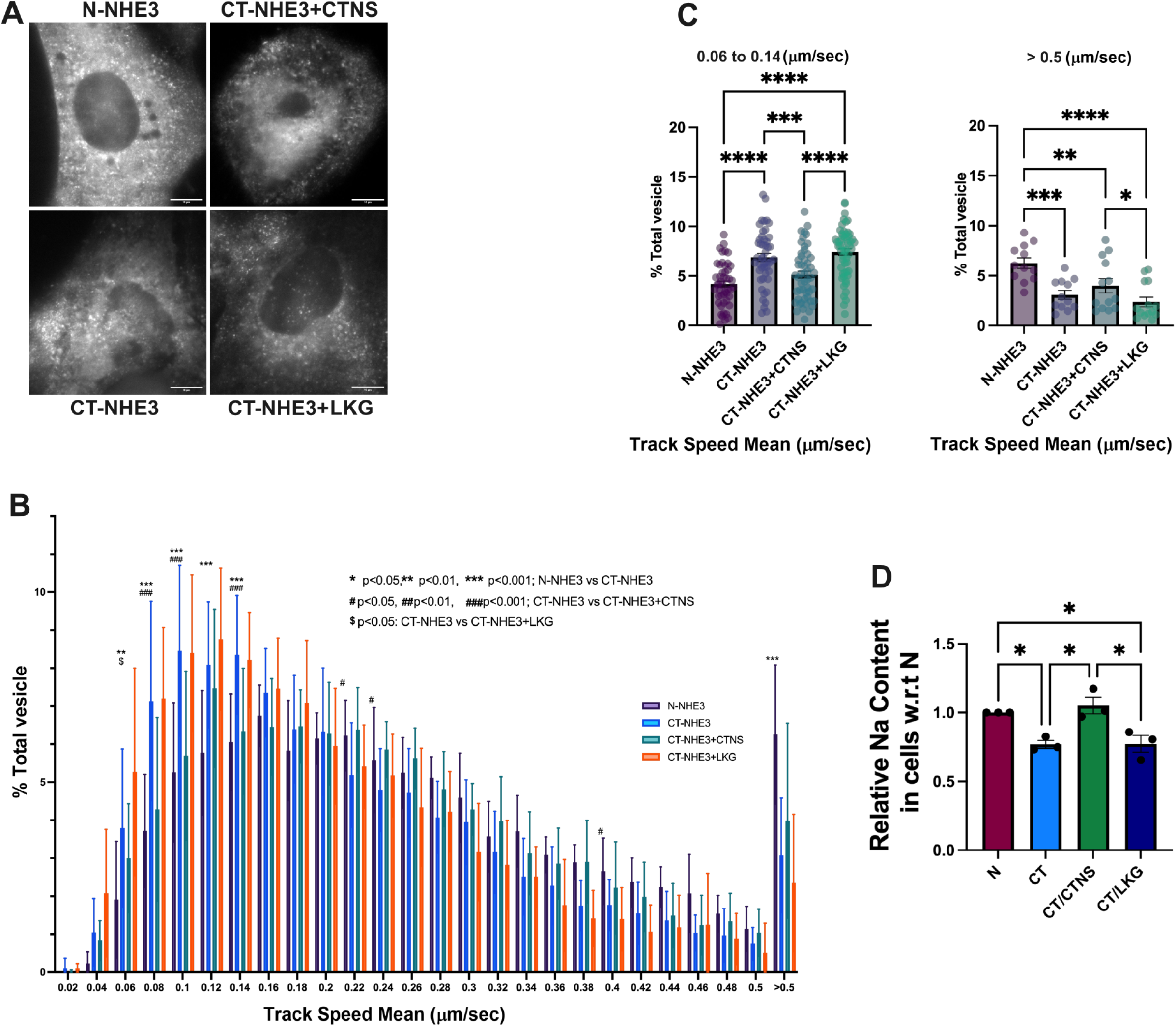
NHE3 trafficking, and sodium uptake are defective in cystinosis patient’s PTCs. (**A**) Representative immunofluorescence images of NHE3-GFP in human normal (N), cystinosis (CT) PTCs and CT cells transduced with LV-CTNS and LV-CTNS-LKG. Scale bar =10 μm. (**B**). Total Internal Reflection Fluorescence microscopy was used to study GFP-tagged NHE3 trafficking in normal (N) and cystinosis patient (CT) PTCs. Two-way ANOVA was used, and the statistics is shown on the figure. Rescue of NHE3 trafficking in CT PTCs was achieved with the transduction with LV-CTNS but not with LV-CTNS-LKG. (**C**) Bar graph showing comparison of slower (0.06-0.14 μm/sec) and faster (>0.5μm/sec) vesicles. (**D**) Intracellular sodium content in the N, CT PTCs and CT transduced with LV-CTNS or LV-CTNS-LKG following treatment with Sodium Green™ Tetraacetate as determined by flow cytometry. All data represents the means and standard error of mean (SEM) from three biological replicates. For (**C** and **D**) One way analysis of variance (ANOVA) was used, \**P* < 0.05; \*\**P* < 0.01; \*\*\**P* < 0.001; \*\*\*\**P* < 0.000.

### *In vivo* study of Nhe3 expression in *Ctns^-/-^* mice and potential rescue by HSPC transplantation

To confirm the *in vitro* results on the impact of cystinosin deficiency on NHE3, we studied Nhe3 expression in the mouse model of cystinosis and assessed whether any defect could be restored following the transplantation of GFP^+^ WT HSPCs. We studied different groups of mice composed of age-matched sex-matched WT, untreated *Ctns*^-/-^ mice, *Ctns*^-/-^ mice transplanted with *Ctns^-/-^* HSPCs (Mock) and *Ctns*^-/-^ mice transplanted with WT GFP^+^ HSPCs (Test) (Fig. 5A). The mice were transplanted at 2 months of age and sacrificed 6 months later. We first investigated *Nhe2* and *Nhe3* expression in the kidney, and no significant change was observed at the transcript level for both genes in any of the groups (Fig. 5B). Nhe3 protein expression was also unchanged between the groups, but interestingly, Nhe2 expression was significantly increased in the *Ctns^-/-^* mouse untreated, Mock, and Test kidneys compared to WT (Fig. 5C) as observed in the HK2 cell lines. Nhe3 protein is expressed in the brush border of the proximal tubular cells (53). We assessed the expression of Nhe3 protein by immunofluorescence in the murine kidneys, using Lotus tetragonolobus lectin (LTL) as a brush border-specific marker. Nhe3 appeared mislocalized in the proximal tubules of the untreated and Mock *Ctns^-/-^* kidneys as demonstrated by Pearson’s correlation coefficient (Fig. 5D). In contrast, significantly improved colocalization between Nhe3 and LTL was observed in the Test group compared to untreated and Mock *Ctns^-/-^* mice (Fig. 5D and Fig. S4). These results provide *in vivo* evidence that cystinosin deficiency causes Nhe3 mislocalization, and this can be rescued by transplantation of *Ctns*-expressing HSPCs.

**Figure. 5.**
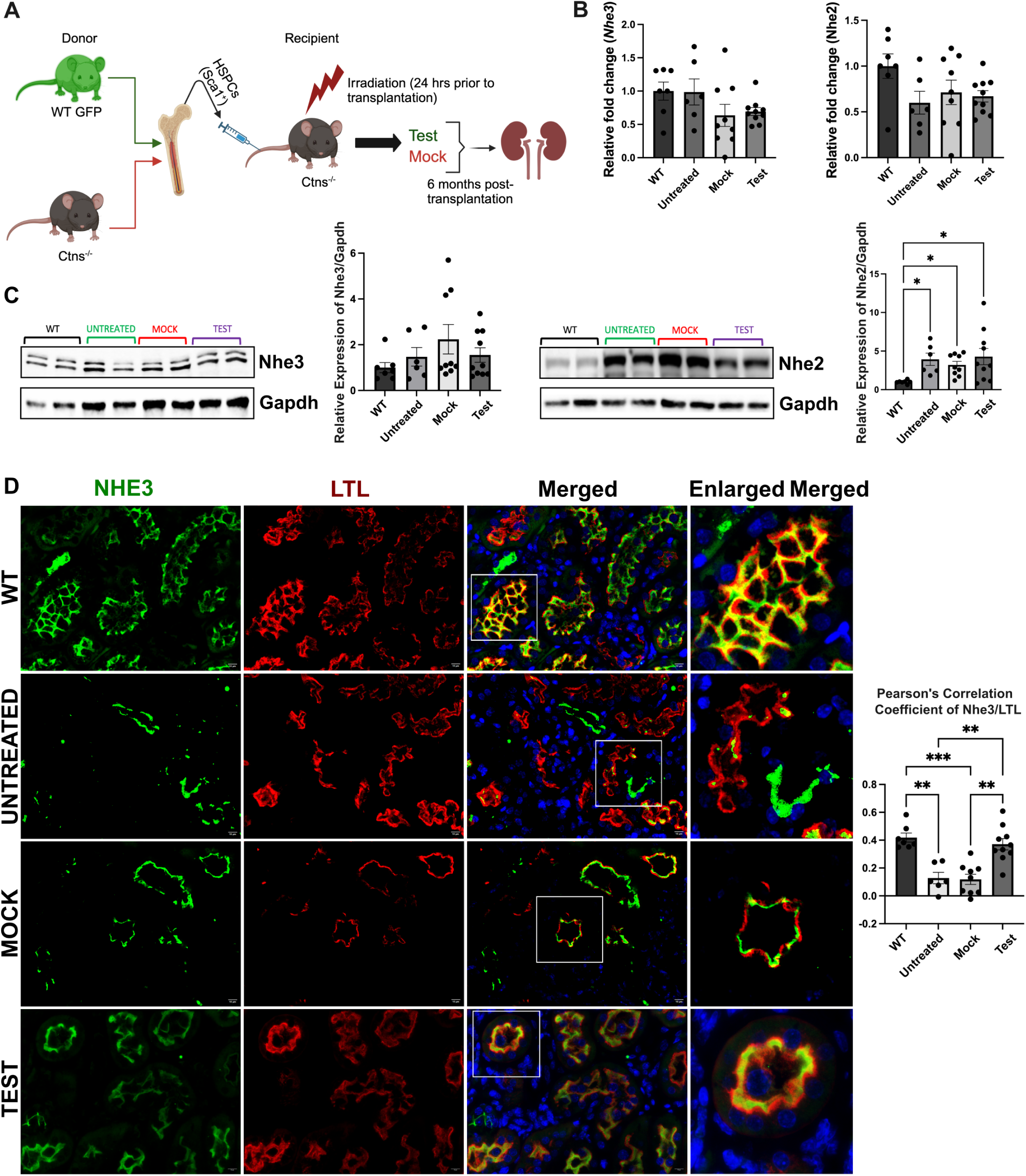
NHE3 localization, but not expression, is affected in *Ctns^-/-^* murine kidneys, and improves/normalizes following WT HSPC transplantation. (**A**) Schematic representation of WT HSPC transplantation procedure. (**B**) *NHE2* and *NHE3* expression measured by quantitative PCR in WT mice, untreated *Ctns^-/-^* mice (untreated), *Ctns^-/-^* mice transplanted with *Ctns^-/-^* HSPCs (Mock), and *Ctns^-/-^* mice transplanted with WT HSPCs (Test mice). (**C**) Representative western blot analysis and quantitative analysis of Nhe3 and Nhe2 expression with GAPDH used as a loading control. (**D**) Representative immunofluorescence images of kidney sections from WT, Ctns^-/-^, Mock and Test mice, stained with anti-NHE3 (green) and anti-Lotus tetragonolobus lectin (LTL-Rhodamine conjugated) (red). Corresponding co-localization quantification between NHE3 and LTL was performed using Pearson Correlation Coefficient. Scale bar = 10μm. Bar graphs are presented as the mean ± SEM. One way analysis of variance (ANOVA), \**P* < 0.05; \*\**P* < 0.01; \*\**P* < 0.01. For data in figure (B-D) each dot represents an independent biological replicate.

### Human cystinosis patient kidney exhibit mislocalized NHE3 expression

We verified the impact of the absence of cystinosin on NHE3 localization in human kidney. Formalin-fixed paraffin-embedded kidney biopsy tissue was obtained from a patient affected with cystinosis and donor control, and sections were stained for NHE3 and LTL marker. In the normal kidney sample, we observed a high degree of colocalization between NHE3 and LTL as demonstrated by Pearson’s correlation coefficient (Fig. 6). However, in the patient tissue, NHE3 and LTL co-localization was significantly decreased (Fig. 6). This data confirms that NHE3 is mislocalized in the kidneys of patients affected with cystinosis.

**Figure. 6.**
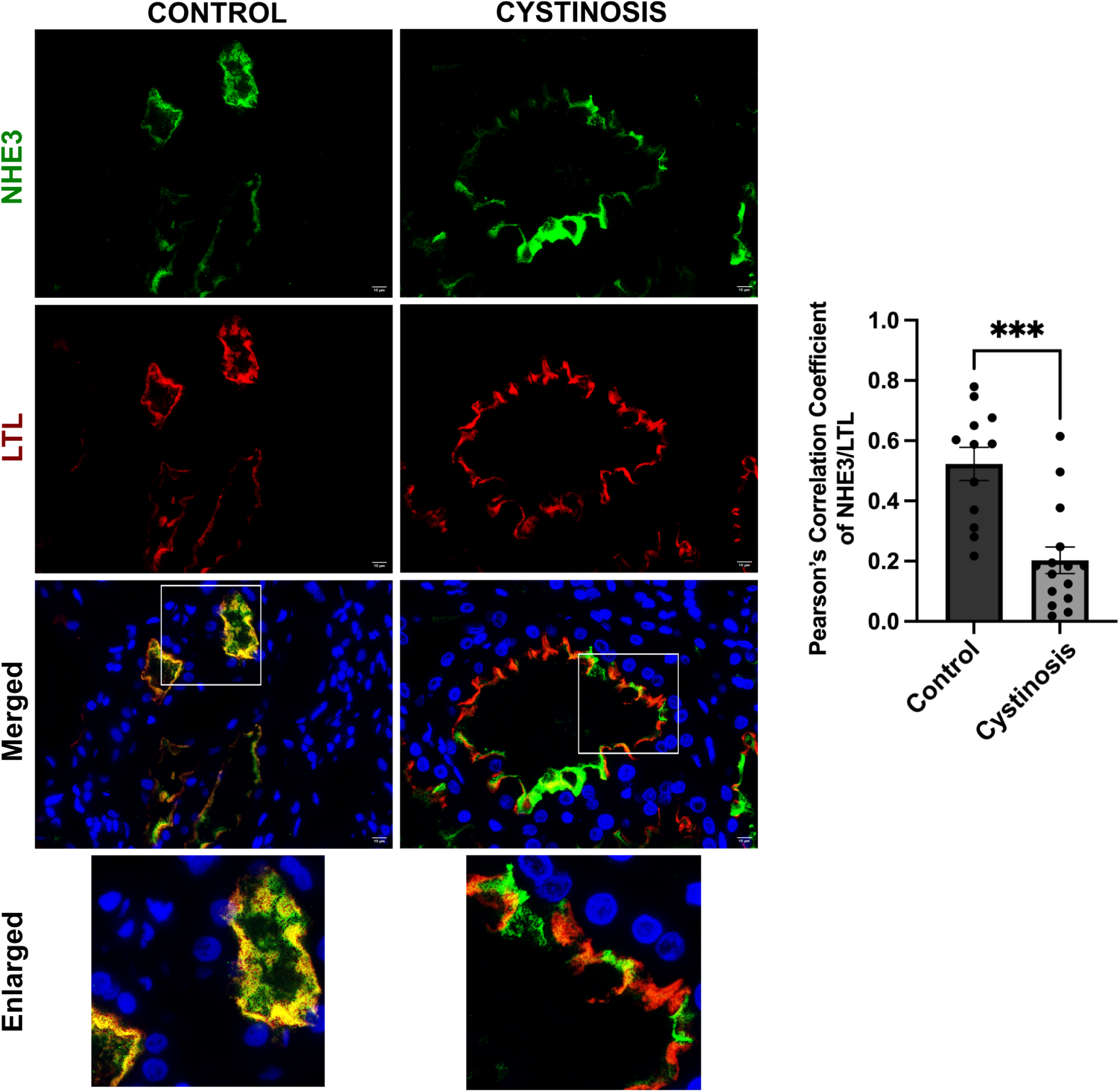
NHE3 localization in human control donor and cystinosis patient kidney tissue sample. Representative immunofluorescence images of formalin fixed paraffin embedded (FFPE) human kidney sections from a control donor and a cystinosis patient. Sections were stained with anti-NHE3 (green), anti-Lotus tetragonolobus lectin (LTL-Rhodamine conjugated) (red) antibodies and DAPI (nuclei). Scale bars = 10 μm. The quantitative data of colocalization between NHE3 and LTL was determined by Pearson Correlation Coefficient. The data are presented as the mean ± SEM. Two-tailed Student’s t-test was used; \*\*\**P* < 0.001. The data in the figure represents the means and standard error of mean (SEM) from three biological replicates.

## Discussion

In this study, we report a novel interaction between cystinosin and a member of the sodium/hydrogen exchanger, NHE3, which plays a pivotal role in proximal tubular cell homeostasis. NHE3 is located at the apical membrane of kidney proximal tubules and contributes to the majority of renal sodium absorption (26). In addition, NHE3 also indirectly contributes to bicarbonates reabsorption which is essential for maintaining acid base homeostasis and overall kidney function (27). Furthermore, NHE3 undergoes dynamic cycling between the apical plasma membrane and the early endosomal compartment, participating in albumin endocytosis in proximal tubules (46, 49). The Na^++^/H^++^ exchange regulatory cofactor (NHERF) family are involved in NHE3 regulation, in multiple aspects of the regulated trafficking of NHE3 (54–57). However, the mechanism by which NHERF proteins regulate the balance of NHE3 between the plasma membrane and endosomes has remained unclear, since the NHERFs are primarily localized to the plasma membrane, with no identified endosomal localization (58).

The reason underlying the pathogenesis of the renal Fanconi syndrome in cystinosis is still unresolved. Despite ubiquitous expression of cystinosin, its first manifestation is renal Fanconi syndrome before one year of age, revealing early severe dysfunction of the proximal tubular cells and reflecting their unique vulnerability in this disease. Decades of research and successive hypotheses have attempted to explain the pathogenesis of the Fanconi syndrome in cystinosis from global metabolic or redox defects (59–61), to impaired lysosomal trafficking and proteolysis (62–64) and to autophagy impairment (15, 17–19, 65). These studies emphasize the importance of the presence of cystinosin *per se* for cellular homeostasis, that goes beyond its transport function. However, the underlying mechanism causing renal FS in cystinosis has yet to be explained. Cystinosin belongs to PQ-loop protein family which are predicted to function as cargo receptors in vesicle transport (66), potentially linking cystinosin to broader lysosomal or endosomal trafficking processes beyond cystine transport. Our investigation of the interaction of cystinosin with NHE3 in the PTCs revealed that these two proteins co-localized within the endosomes, distinguishing this complex from the lysosomal cystine transporter function of cystinosin. Most importantly, in human *CTNS*-deficient PTCs, NHE3 was mislocalized to the endoplasmic reticulum and NHE3 vesicular trafficking was impaired, providing the first evidence that cystinosin is involved in NHE3 trafficking. These novel findings are conserved in primitive models such as the *P. Pastoris* yeast model underscoring their importance. Indeed, we observed that the yeast orthologues of the NHE proteins and cystinosin, Nhx1 and Ers1, respectively, colocalized in the endocytic vesicles. Furthermore, mislocalization and deficient trafficking of Nhx1 were also observed in the Ers1-deficient lines. Ers1 has also been shown to move from phagosome to lysosome *in C.elegans* (66, 67). In *P. Pastoris,* Δ*ers1S* Δ*ers1L* double mutant, Nhx1 protruded beyond the late endosomes and a large fraction localized into the vacuolar matrix. This suggests that the double deletions of Ers1 (S and L) affect late endosome/multivesicular body (MVB) morphology. Nhx1 is reported to be crucial for MVB vesicle formation in late endosomes (68) and MVB-vacuolar lysosome fusion, a key step in surface protein degradation during endocytosis (69). This correlates with the phenotypic trait we observed under double deletion of ERS1 (S and L), thus suggesting that Ers1 might function to retain Nhx1 in the endocytic pathway through an unknown retrograde/recycling mechanism. In contrast, deletion of Ers1 in *S. cerevisiae* did not lead to cystine accumulation (42), and we obtained similar results in *P. pastoris* strains lacking Ers1S, Ers1L, or both, even under nitrogen starvation conditions. These results suggest that cystinosin’s role in vesicular trafficking may be its primary function rather than serving as a cystine transporter.

This study focused on the human NHE isoforms expressed at the plasma membrane, specifically examining their role in maintaining sodium homeostasis in the kidney such as NHE2 and NHE3. Intracellular isoforms, such as the vacuolar and endosomal members (e.g., NHE6 and NHE8), which are primarily involved in pH and ion balance, were not included in our investigation (70). Surprisingly, while both cystinosin and cystinosinLKG interacted with NHE3, rescue of the expression of *CTNS*, but not *CTNS-LKG,* was able to restore normal trafficking of NHE3 in cystinosis patient PTCs and improve sodium and albumin uptake in CTNS-deficient PTC lines. The specific role of cystinosinLKG is unknown. This isoform was shown to decrease cystine accumulation and apoptosis in Ctns-deficient PTCs (71). In tissues, *CTNS-LKG* mRNA was found to be expressed at a level included between 5-20% of the canonical isoform in most tissues except in testis, in which *CTNS-LKG* mRNA level is similar to *CTNS* (72). CystinosinLKG is localized at both the lysosome and the plasma membranes, with the highest levels observed in cells involved in secretory activities, such as renal tubular epithelial cells, islets of Langerhans, mucoserous glands of the bronchial epithelium, melanocytes, keratinocytes, and Leydig cells. This suggests that this isoform may play a key role in intracellular trafficking and secretory functions (72). In this study, we showed that only cystinosin is involved in NHE3 trafficking, and thus, while both isoforms are involved in cystine transport (22, 73), they have distinct functions for cellular trafficking that could be specific to proteins or cell types. Further studies will be required to determine the specific roles of cystinosin versus cystinosinLKG in each cell type.

We confirmed the abnormal NHE3 localization in the PTCs in the murine *Ctns^-/-^* kidneys and cystinosis patient kidney biopsies. Loss of apical multi-ligand endocytic receptors, megalin, and cubilin, has also been observed in *Ctns^-/-^* mice (62, 74). Under normal physiological conditions, NHE3 specifically interacts with megalin in proximal tubules, participating in megalin/cubilin-mediated endocytosis (46, 49, 75). Megalin and cubilin facilitate receptor-mediated endocytosis of low-molecular-weight proteins in the kidney and are expressed at the renal brush border (76–78). Therefore, abnormal NHE3 trafficking to the brush border in cystinosis models could also help explain the loss of megalin and cubilin, accounting for the overall water, small proteins and solute loss associated with the renal FS. In addition, several reports suggested absorptive functions for both NHE3 and NHE2 isoforms in intestinal and renal proximal tubule brush border, with NHE3 predominantly assuming the major role (24, 26, 79). We observed an increased expression of NHE2 in *CTNS*-deficient PTCs and in *Ctns^-/-^* kidney, probably acting as a compensatory mechanism to the loss of NHE3 at the apical membrane. However, the contribution of NHE2 to absorption, if any, is not significant enough to compensate for NHE3 loss (80, 81).

The current therapy for cystinosis, cysteamine, which allows lysosomal cystine clearance, delays disease progression but fails to correct the Fanconi syndrome. Some recent studies indicate that starting cysteamine treatment immediately after birth, particularly in asymptomatic infants, may slow the progression of renal Fanconi syndrome. However, these observations are largely anecdotal and require further investigation to determine the optimal timing for treatment initiation, overall drug tolerance, compliance, and the long-term effectiveness of the therapy in mitigating the severity of Fanconi syndrome and kidney dysfunction (82, 83). Our study supports that the renal FS in cystinosis is due to cystinosin-specific function in the PTCs, and not due to the toxic impact of the cystine storage. Therefore, the treatment of the FS in cystinosis requires the restoration of cystinosin in the PTCs. We previously demonstrated that stable engraftment of bone marrow-derived cells from *Ctns*-expressing HSPC transplantation in *Ctns^-/-^* mice resulted in a reduction in cystine content in tissues and long-term preservation of the kidney, and improvement of the renal FS (8, 9). The primary mechanism underlying this therapeutic effect involved the formation of TNTs by HSPC-derived macrophages, facilitating the transfer of healthy lysosomes containing functional cystinosin to the diseased host cells, including the proximal tubular cells (10, 84). In this study, our results showed that NHE3 localization at the brush border was significantly improved in the *Ctns^-/-^* mice transplanted with WT HSPCs, further supporting restoration of cystinosin protein in the PTCs of the treated mice. The knowledge obtained from this study will also be pertinent to the ongoing stem cell gene therapy clinical trial for cystinosis as well as other rare kidney diseases (85).

Renal Fanconi syndrome is a multifactorial condition characterized by generalized dysfunction of the proximal tubules, leading to impaired reabsorption of various substances. While NHE3 plays a role in sodium and bicarbonate reabsorption, its dysfunction alone may not fully explain the spectrum of FS pathology. Indeed, mice and patients with genetic loss-of-function mutations in *NHE3*, no Fanconi syndrome has been described (86, 87) supporting the fact that cystinosin mediated FS is a multifactorial implication rather than a single gene effect. As such, novel roles of cystinosin in PTCs such as its involvement in autophagy (15–19), TFEB expression (20), and inflammation (21) has been identified. In addition, loss of expression and mislocalization of proteins like Sodium-Glucose Cotransporter 2 (*SGLT2*), megalin (*LRP2*), and Type IIa Sodium-Dependent Phosphate Transporter (*NPT2*) have been reported, though the mechanisms have not been fully explored (74). Interestingly, NHE3 interacts directly or indirectly with these proteins (75, 88–90). Restoring cystinosin function may alleviate these effects by stabilizing NHE3 localization and function, potentially addressing one of the primary causes of FS. However, while the restoration of NHE3 expression following *CTNS* supplementation is an intriguing finding, it does not prove that NHE3 dysfunction is the sole factor contributing to the development of FS in cystinosis. The relationship may be indirect or part of a broader network of cellular changes.

In summary, cystinosin interacts with NHE3 and plays a crucial role in its expression, trafficking, and function. Given that NHE3 has an essential role in the PTCs, interacting directly and indirectly with several transporters, this novel finding provides a mechanism underlying the renal Fanconi syndrome in cystinosis. These findings may lead to novel therapeutic approaches for cystinosis and provide valuable insights into the current HSPC gene therapy clinical trial for cystinosis.

## Methods

### Mice

All mice used in this study were of the C57BL/6 background. The C57BL/6 *Ctns^-/-^* transgenic mouse model was generously provided by Dr. Corinne Antignac (Inserm U983, Paris, France) (6, 7). Genotyping of the mice was conducted using standard PCR reactions with primers detailed in Table S3. For hematopoietic stem/progenitor cell transplantation experiments, GFP transgenic mice expressing enhanced GFP cDNA under the control of the chicken beta-actin promoter (C57Bl/6-Tg (CAG-EGFP)1Osb/J; 003291, Jackson Laboratory) were utilized as donors. The mice were housed in a controlled environment with regulated temperature and humidity, maintained on a 12-hour light/dark cycle, and provided free access to water and food. Both male and female mice were included in the experiments. Each experimental and control group comprised 6–10 mice, analyzed at approximately 8 months of age; details regarding sex and group numbers are provided in the results section. All mice were bred at the University of California, San Diego (UCSD) vivarium, and all experimental procedures were conducted in accordance with protocols approved by the UCSD Institutional Animal Care and Use Committee.

### Mammalian cells

Human N and CT PTCs were generously provided by Dr. Corinne Antignac. They are transformed PTCs isolated from the urine of a healthy donor (N), and from a cystinosis patient (CT) that has a 57kb deletion in the heterozygous state and a G>T transition at the donor splice site of exon 7 (800G>T) (91). HK-2 (proximal tubule cells) and *CTNS^-/-^* HK-2 cells were provided by Dr. Sergio Catz (45). The human cell line HEK293T (embryonic kidney) was obtained from ATCC and used for producing lentiviral particles for LV-CTNS-DsRed; LV-CTNS-LKG-DsRed, LV-NHE1-GFP, LV-NHE2-GFP, and LV-NHE3-GFP, and for the immunoprecipitation studies.

### Yeast cells

The studies were carried out using methylotrophic yeast *Pichia pastoris*.

### Human kidney biopsy

Kidney biopsy from a patient affected with cystinosis was obtained and the studies have been approved by the institutional review board committee (IRB #141150) and have been performed in accordance with the ethical standards. Patient signed informed consent. The biopsy was provided as formalin-fixed paraffin-embedded (FFPE) blocks. Tissue from non-cystinosis healthy donor was obtained from Zyagen (HP-901). 5 μm sections were cut and stained by rabbit/rat anti-NHE3 (1:50; BiCell Scientific), LTL Rhodamine conjugate (1:100; BiCell Scientific) and DAPI stain.

### Statistical analysis

The statistically significant differences were determined using GraphPad software Prism v.10.0. All data are presented as mean ± SEM and *p* values less than or equal to 0.05 were considered statistically significant. Comparisons between 2 groups were performed using an unpaired or paired 2-tailed Student’s t test. One-way or two-way analysis of variance (ANOVA) was used to compare multiple groups. For the mice studies, all cohorts in the experiments were age-matched and sex-balanced. Experimenters were blinded to the genotype of the samples as much as possible. We did not exclude any animals from our experiments.

The details of material and methods are listed in the Supplementary Methods and Supplementary Tables S1–S4.

## Acknowledgements

The study was supported by funds from the Cystinosis Research Foundation, the California Institute of Regenerative Medicine (CIRM, CLIN2-11478 and TRAN1-13983), the National Institute of Health (NIH) R01-NS108965, R01NS135162, R01-AG086443 to S.C; R01DK110162, and P01HL152958 to SDCatz, the MPS Society, and the Friedreich’s Ataxia Research Alliance (FARA). The UCSD Neuroscience Microscopy Shared Facility was funded by grant P30-NS047101. The Automated Zeiss LSM 980 Airyscan 2 Multiscale super-resolution Confocal microscope was funded by an NIH S10 grant S10OD030417 to SDCatz. We gratefully acknowledge Dr. Corinne Antignac for generously providing the mouse model of cystinosis and sharing the human N and CT PTCs and Dr. Benjamin Glick for providing the yeast strain PPY12+Sec7-DsRed. We acknowledge the cystinosis patient for donation of the kidney tissue. We acknowledge Jay Sharma, and the students Samantha Diaz, Pruthva Mania, Andrea Kelly, Tariq Aniff, Nia Asbill and Junmyung Lee for their help on the project.

## Data, Materials, and Software Availability

All the data that support the findings of the study are available from the corresponding author upon reasonable request.

## Author contributions

SC, VK, JCF, CR, and SDCatz contributed to the conception of the study. VK, JCF, CR, MAK, CT, XM, KB, IM, RABG, AS, RC performed the experiments. VK, JCF, MAK, SDCatz, SC analyzed and interpreted the data. SC, JCF, and VK wrote the manuscript. All authors discussed the results and approved the final version of the manuscript.

## Figure Legends

**Figure S1. - Sequence alignments and AlphaFold-predicted structures of cystinosin protein isoforms in humans, *P. pastoris*, and *S. cerevisiae***. **(A)** Clustal Omega multiple sequence alignment of human cystinosin (Uniprot accession number A0A0S2Z3K3), cystinosinLKG (A0A0S2Z3I9), *P. pastoris* Ers1S (C4R5N7), *P. pastoris* Ers1L (C4R120), and *S. cerevisiae* Ers1 (P1761). Black boxes indicate amino acid identity, grey boxes indicate amino acid similarity, with the red-boxed area highlighting the comparison of the seven-transmembrane domains of cystinosin across the different organisms. We used BoxShade for highlighting (https://junli.netlify.app/apps/boxshade/#forms::boxshade). The multiple sequence alignment was performed using Clustal Omega from EMBL (https://www.ebi.ac.uk/jdispatcher/msa/clustalo). The threshold for comparison is set by default to greater than 50%. **(B)** Displays images of the AlphaFold-predicted structures of the various cystinosin isoforms in each organism described in **(A).**

**Figure S2. - Cystinosin does not interact with NHE1.** Immunoprecipitation assay performed in HEK293T cells stably expressing NHE1, cystinosin, and cystinosinLKG showing no interaction between NHE1 and both cystinosin and cystinosinLKG. Input lysates show proper expression of the proteins with glyceraldehyde-3-phosphate dehydrogenase (GAPDH) used as a loading control. Pull down with anti-IgG was used as a negative control.

**Figure S3. - Cystinosin deficiency impairs NHE3 trafficking in human WT and *CTNS^-/-^* HK2 proximal tubular cells.** (**A & B)** Representative immunofluorescence images of NHE3-GFP (green) with markers of lysosome (LAMP1) (red) (**A**) and Golgi (GM130) (red) (**B**) with their corresponding quantification showing defect in NHE3 subcellular localization in *CTNS^-/-^* HK2 cells. Scale bars = 20 μm Bar graphs are presented as the mean ± SEM. \**P* < 0.05; \*\*\**P* < 0.001 using two-tailed Student’s t test.

**Figure S4. - GFP^+^ WT HSPC distribution in the kidney following WT HSPC transplantation.** Overview of immunofluorescence images of entire kidney section from the Test mice stained with anti-GFP (green), anti-NHE3 (magenta) and anti-LTL (red) antibodies. This image shows the distribution of the GFP^+^ WT HSPC-derived cells as well as the colocalization of LTL and NHE3 within the kidney of *Ctns*^-/-^ mice transplanted with WT HSPCs. Scale bars = 500μm.

## Supplementary Tables

**Table S1.**
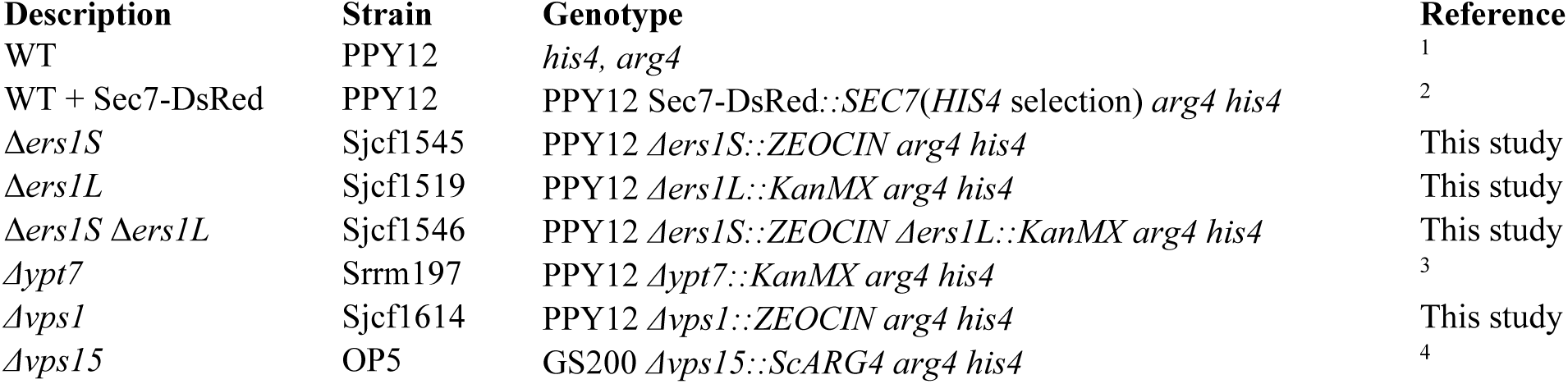
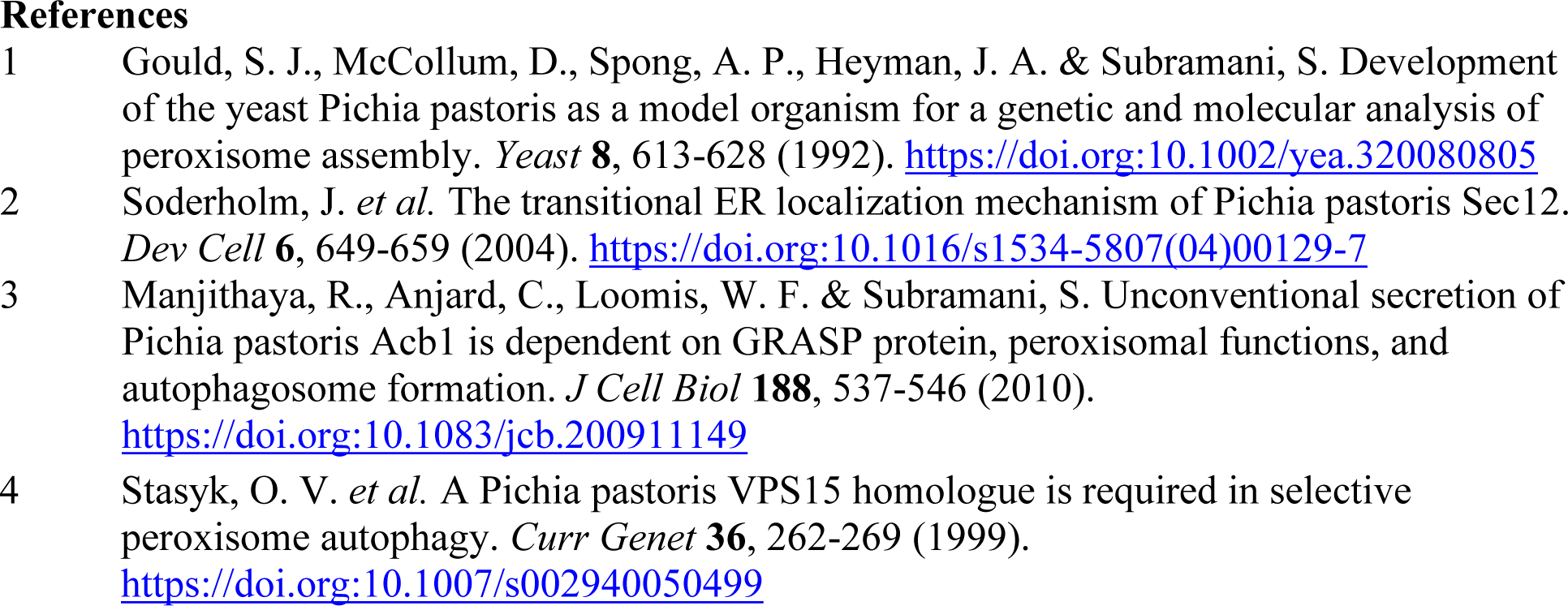
*Pichia pastoris* strains.

**Table S2.** *Pichia pastoris* plasmids.

| Plasmid | Promoter | Fusion Protein | Integration Locus | Selectable Marker |
| --- | --- | --- | --- | --- |
| pJCF728 | *ERS1S* | Ers1S-GFP+ | *ARG4* | *ARG4* |
| pJCF155 | *NHX1* | Nhx1-GFP | *ARG4* | *ARG4* |
| pTA12 | *VPS8* | Vps8-2xmCherry | *VPS8* | *NAT* |
| pTA8 | *NHX1 / ERS1S* | Nhx1-VC / Ers1S-VN | *ARG4* | *ARG4* |
| pTA3 | *NHX1* | Nhx1-VC | *HIS4* | *HIS4* |
| pTA2 | *ERS1S* | Ers1S VN | *HIS4* | *HIS4* |

**Table S3.**
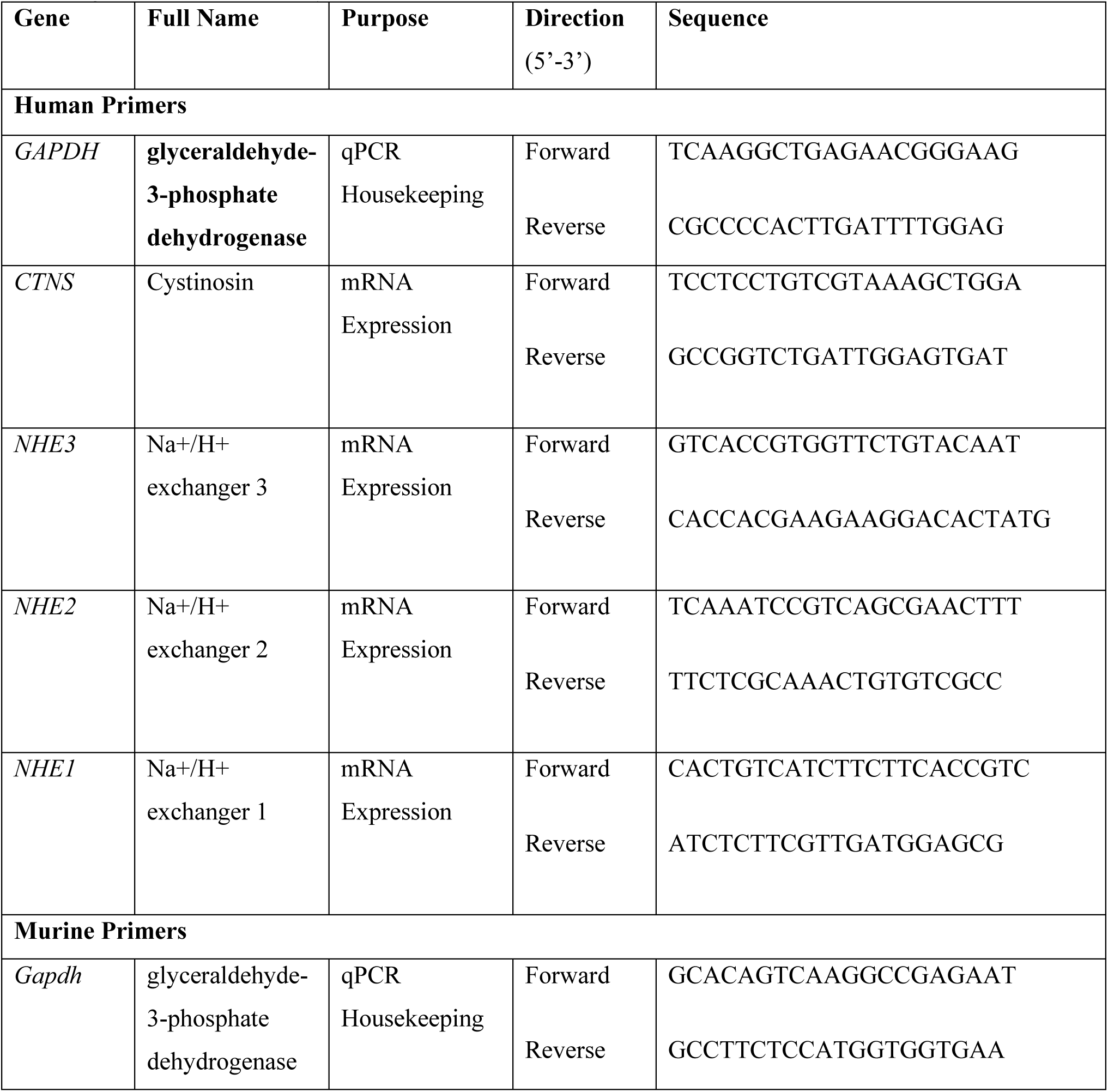
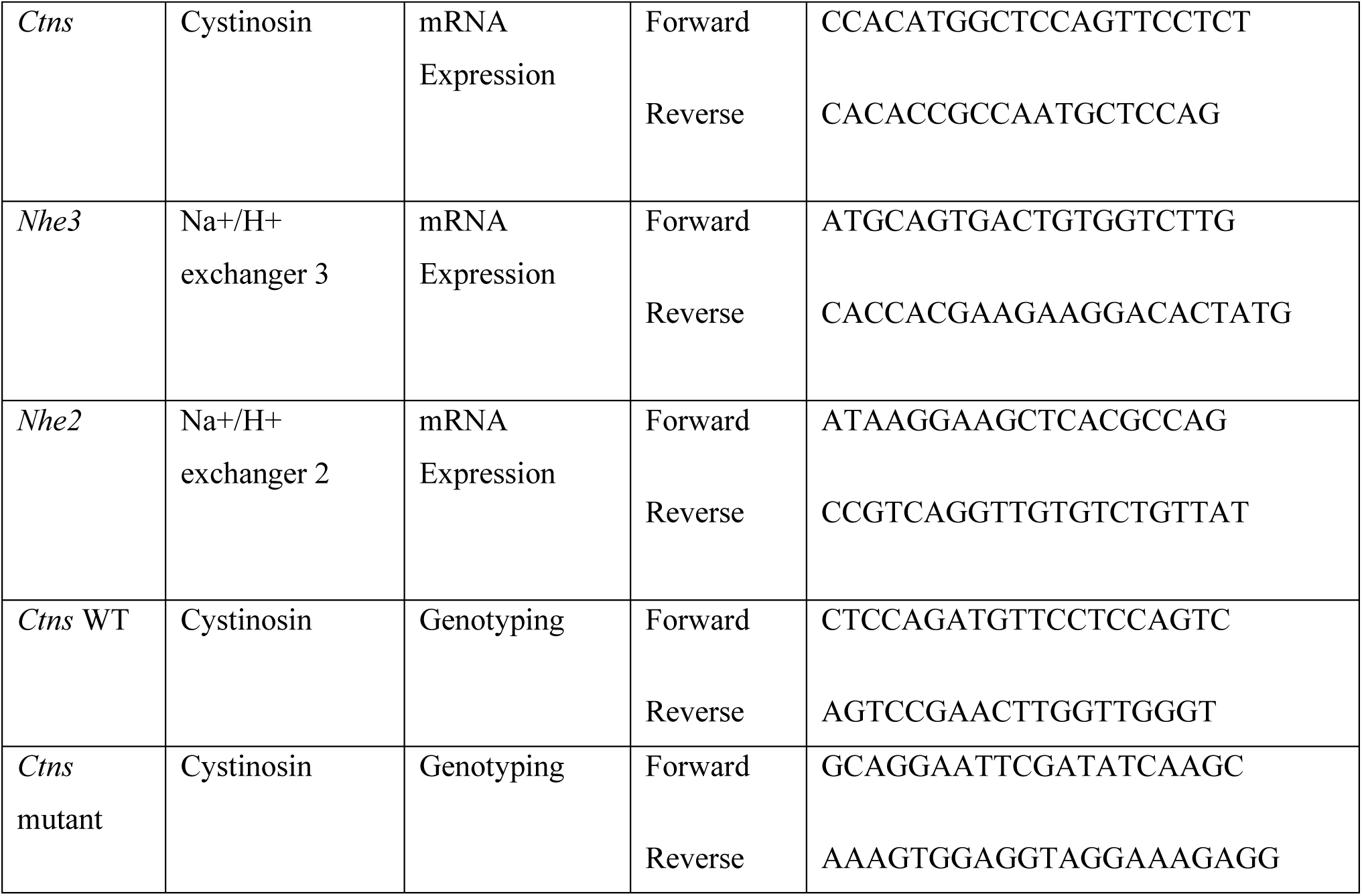
Primer sequences for qPCR. All primers were reconstituted at 100 μM and used at a working concentration of 5 μM.

**Table S4.**
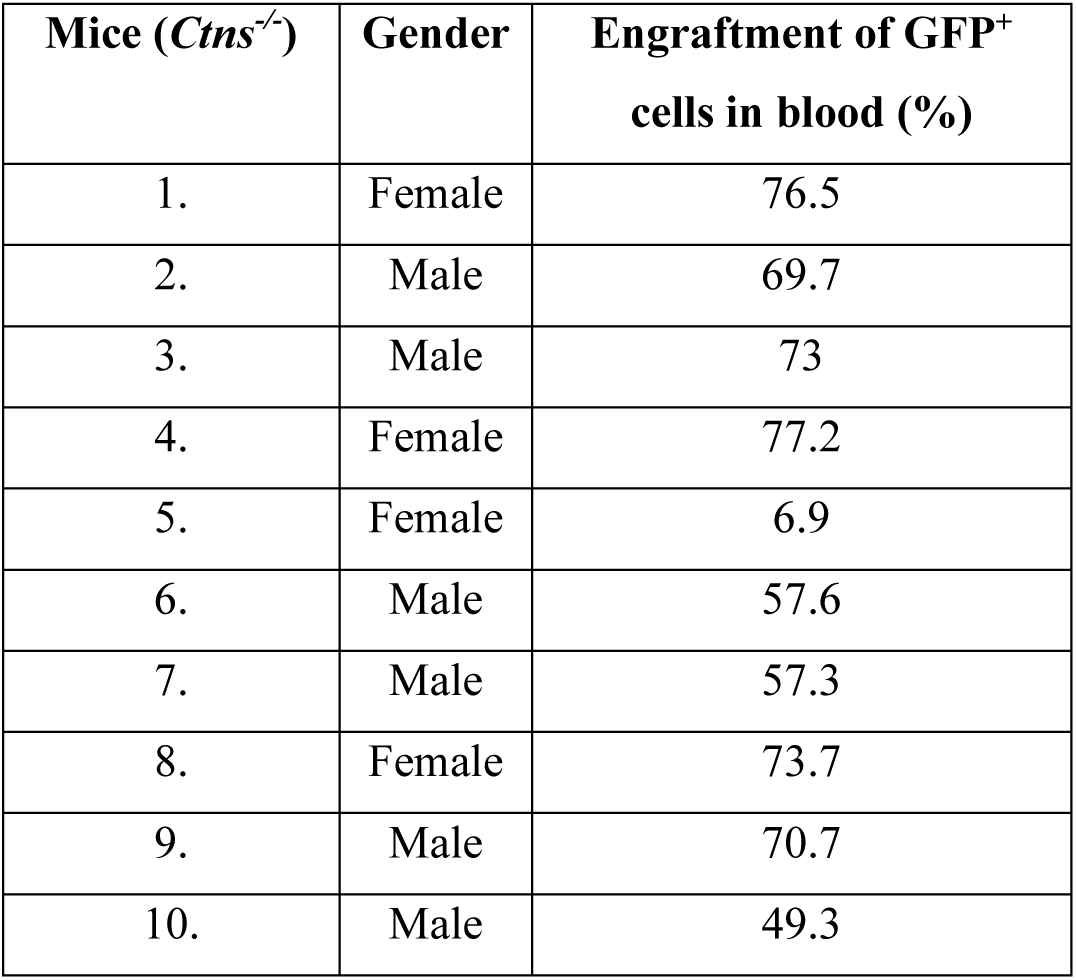
Donor-derived HSPC engraftment in *Ctns^-/-^* mice transplanted with GFP^+^ HSPCs.

| Mice (*Ctns*<sup>-/-</sup>) | Gender | Engraftment of GFP<sup>+</sup> cells in blood (%) |
| --- | --- | --- |
| 1. | Female | 76.5 |
| 2. | Male | 69.7 |
| 3. | Male | 73 |
| 4. | Female | 77.2 |
| 5. | Female | 6.9 |
| 6. | Male | 57.6 |
| 7. | Male | 57.3 |
| 8. | Female | 73.7 |
| 9. | Male | 70.7 |
| 10. | Male | 49.3 |

## SUPPLEMENTARY METHODS

### Lentiviral particle production

HEK293T cells line was cultured in DMEM supplemented with 10% heat inactivated FBS, with a prescribed dose of Penicillin/Streptomycin (Invitrogen, Life Technologies) added as recommended antibiotics. All cells were maintained at 37°C in a humid incubator with 5% CO2. Lentiviral particles were produced using the 3 packaging plasmids (pMDLg/pRRE, pHCMV-g, pRSV-Rev) (12) along with the plasmid carrying *CTNS-DsRed*, *CTNS-LKG-DsRed,* and *NHE1-GFP*, *NHE2-GFP*, *NHE3-GFP* and were transfected using calcium phosphate protocol (12). Vector particles were harvested in media and concentrated through ultracentrifugation at 25,000 rpm for 2 hours at 4°C. The titers of the concentrated virus, finally dissolved in Stemspan medium (StemCell Technologies, Vancouver, British Columbia, Canada), were determined by infecting HEK293T cells with serial dilutions of the virus preparations and evaluating them via flow cytometry and droplet digital (dd) PCR. Subsequently, the virus was introduced to HEK293T and maintained for 14 days to establish the stable cell lines. The GFP and DsRed positive cells were selected using fluorescence-activated cell sorting (FACS).

### Yeast studies

#### Strains and plasmids used are shown in Supplementary Table S1 and S2, respectively

Medium used in this study: YPD (2% glucose, 2% bacto-peptone, 1% yeast extract), YNB (0.17% yeast nitrogen base without amino acids and 0.5% ammonium sulfate), YNB-N (0.17% yeast nitrogen base without amino acids and ammonium sulfate), CSM (complete synthetic medium of amino acids and supplements), glucose medium (2x YNB, 0.79g/L CSM, 0.04 mg/L biotin, 2% dextrose), SD-N or starvation medium (1x YNB-N and 2% dextrose)

#### Extraction of intracellular cystine

Samples (50 ml) of cultures were harvested by centrifugation at 2,000 g and washed twice in 100 mM phosphate buffer (pH 7.5). The cells were suspended in 5% trichloroacetic acid (TCA) in water and broken with 0.5-mm glass beads. Cell debris was removed by centrifugation (10,000 g) at 4°C, and the supernatant was stored at -20°C for MS analysis of cystine content.

#### Fluorescence Microscopy

Cells were grown in YPD at 30°C until exponential phase (1–2 OD600/mL), washed twice with sterile water, and then transferred to glucose medium for 6 hours. Mid-log cells were then pelleted, 1.5 μl of cells was mixed with 1% low melting point agarose and placed on a glass slide with a coverslip and imaged using 63× or 100× magnification on a Carl Zeiss Axioskop 2 MOT microscope (Carl Zeiss Microscopy, Gottingen, Germany). Images were taken on an AxioCam HRm digital camera (Carl Zeiss MicroImaging GmbH, Gottingen, Germany); no digital gain was used, exposure was adjusted as needed, except for BiFC which was kept constant during microscopy in different strains. Images were processed using AxioVision software V4.8.2.0 (Carl Zeiss Microscopy, White Plains, NY, USA). The images are representative results from experiments conducted at least in triplicate. Methodology to determine late endosome (LE) or trans-Golgi network/early endosome (TGN/EE) localization in different background strains is as follows: Cells showing distinct LE labeling with Vps8-2xmCherry or TGN/EE labeling with Sec7-DsRed were first marked in the red channel in the AxioVision software. Then, these marks were analyzed for colocalization in the green channel with the GFP-tagged proteins or fluorescence obtained from Bimolecular Fluorescence Complementation (BiFC). FM4-64 staining was visualized after a short pulse (∼3 min) followed by a chase in the presence of the quencher (4-Sulfonato calix [8] arene, sodium salt) SCAS (BIOTIUM cat#70037).

### Immunoprecipitation (IP) assay

HEK293T stably transduced with various combinations of LV-CTNS-DsRed; LV-CTNS-LKG-DsRed, LV-NHE1-GFP, LV-NHE2-GFP, and LV-NHE3-GFP, were cultured and harvested from 15 cm dish plates. Whole cell extracts were prepared using IP Lysis Buffer (50 mM Tris-HCL pH 7.5, 150 mM NaCl, 1mM EDTA, 1% Igepal CA-630, 10% Glycerol, 0.5 mM DTT) supplemented with freshly added PIC, PMSF & DTT. Supernatants were collected by centrifugation at 13000 x g for 10 min at 4°C. Protein conc. was measured by Pierce BCA Protein Assay Kit. 10% Input was kept at -80°C for later use. Prior to immunoprecipitation (IP), lysates underwent pre-clearing via incubation with Dynabeads at 4°C for 30 mins. GFP pull-down was executed using GFP-Trap Magnetic Beads (Chromotek-gtma), while DsRed pull-down utilized Protein A Dynabeads pre-coupled with RFP Antibody (Chromotek-6G6) at 4°C for 2-4 hrs. The pre-coupled beads were then incubated with the 500ug-1mg protein lysate at 4°C O/N. Next day, the tubes were replaced, and the beads were washed with IP Lysis Buffer followed by IP Wash Buffer (50mM Tris-HCL pH 7.5, 200mM NaCl, 1mM EDTA, 1% Igepal CA-630, 10% Glycerol,1mM DTT) (with freshly added PIC, PMSF & DTT). Elution of proteins was achieved using 30 μl of 2X protein loading dye. Co-immunoprecipitated proteins were resolved by SDS-PAGE and probed for GFP (Abcam, ab290) and DsRed (Chromotek-6G6). Pulldown using normal mouse IgG (sc-2025) or normal rabbit IgG (CST-2729) was used as a negative control.

### Immunoblotting

Cells and murine kidney tissues were homogenized in Pierce RIPA buffer (Sigma, St Louis, MO) supplemented with Protease Inhibitor Cocktail (Sigma, St Louis, MO). Protein concentrations were determined using the Pierce BCA Protein Assay Kit and 30-50μg of proteins were loaded onto SDS-PAGE gels for subsequent immunoblotting following standard protocols. The proteins transferred onto the PVDF membrane was incubated with the primary antibodies at a 1:1000 dilutions in 5% BSA in TBS-T, with overnight incubation at 4°C for primary antibodies and 1-hour incubation at room temperature for secondary antibodies. The primary antibodies used were rabbit anti-NHE3 (Millipore, AB3085), rabbit anti-NHE2 antibody (Novus, NBP2-38236), mouse anti-NHE1 antibody (Proteintech, 67363-1-1g) and mouse anti-GAPDH antibody (Proteintech, HRP-60004) followed by goat anti-mouse or anti-rabbit horseradish peroxidase-conjugated secondary antibodies. Blots were developed using Enhanced Chemiluminescence (ECL) reagent according to the manufacturer’s protocol (GE Healthcare, Pittsburgh, USA) and imaged using the Azure 600 imager (Azure Biosystems, Dublin, CA, USA). Quantification of bands was carried out using GelQuant.Net software.

### RNA extraction and real-time quantitative PCR (RT-qPCR)

Total RNA was extracted from both cells and mice kidney (after homogenization in Precellys 24) employing the RNeasy Mini Kit as per the manufacturer’s protocol (Qiagen, Hilden, Germany, cat. 74104). Subsequently, 500 ng of RNA was transcribed into cDNA using iScript cDNA Synthesis Kit (Bio-Rad, Hercules, CA, cat. 1708840). For the RT-qPCR reaction setup, 5 μl of iTaq Universal SYBR Green Supermix (Bio-Rad, Hercules, CA, cat.1725121), 3 μl of 1:10 diluted cDNA (2.5 ng/μl), and 1 μl of forward and reverse primer (5 μM each) were combined. The reaction was performed on a CFX96 thermocycler (Bio-Rad) under the following conditions: 95°C (30 s); 40 cycles of 95°C (5 s) and 60°C (30 s); then 65°C (5 s); and 95°C (5 s). Gene expression was quantified utilizing the ΔΔCt method relative to the wild-type (WT) and normalized to the endogenous control (GAPDH). All primer sequences are shown in Table S3.

### Determination of Intracellular Sodium

To measure intracellular sodium concentration [Na^+^], we used the sodium ion fluorescence indicator Sodium Green™ Tetraacetate (Invitrogen, Catalog #S6901) (92). We seeded 0.6 × 10^6^ cells in 24-well plates overnight. The next day, the cells were washed twice with PBS and incubated with Sodium Green™ Tetraacetate at a final concentration of 2μM in dimethyl sulfoxide for 8 minutes. After incubation, the cells were washed three times with PBS to remove excess dye and processed for flow cytometry (BD Accuri C6, BD Biosciences). AAD-7 was used to exclude dead cells. All flow cytometric analyses were performed using BD software. We also conducted this assay following cells treated with NHE3 inhibitor EIPA (5-(*N*-ethyl-*N*-isopropyl)-amiloride (Sigma) at a concentration of 100 μM for 4 hrs. DMSO was used as the vehicle control for EIPA treatment with a final concentration of 0.1%.

### Albumin Uptake Assay

To assess the cells’ ability to uptake albumin, 0.6 × 10^6^ cells were seeded in a 24-well plate overnight. The next day, the cells were washed twice with PBS and incubated with 50 μg/mL Alexa Fluor 555-conjugated albumin (Invitrogen, Cat #A34786) for approximately 16 hours for complete intracellular uptake and processing of albumin. After incubation, the cells were washed three times with PBS to remove excess dye and processed for flow cytometry (BD Accuri C6, BD Biosciences). AnnexinV (Invitrogen, Cat #331200) & 7-amino-actinomycin D (7-AAD) (Invitrogen, Cat # 00-6993-50) was used to exclude dead cells. All flow cytometric analyses were conducted using BD software. We also conducted this assay following cells treated with NHE3 inhibitor EIPA (5-(*N*-ethyl-*N*-isopropyl)-amiloride (Sigma) at a concentration of 100 μM for 4 hrs.

### Total internal reflection fluorescence microscopy (TIRFM) and data analysis

For live-cell TIRFM imaging, human PTCs were seeded in 8-well plates with coverglass bottoms (Lab-Tek borosilicate, Nunc, Thermo) in phenol red-free RPMI medium. Cells were placed on a prewarmed microscope stage, and imaging was done using a 100× 1.45 numerical aperture (NA) TIRF objective on a custom-modified Nikon TE2000U microscope with TIRF illumination. Laser illumination (488 and 543 nm) was angled to create an evanescent field depth of <100 nm. Images were captured on a 14-bit cooled CCD camera (Hamamatsu) controlled by NIS-Elements software (Nikon) at 1-second intervals with 200–600 ms exposure times. Analysis was performed using ImageJ (version 1.43) and Imaris (version 7.0, Bitplane Scientific Software). Granule movement was tracked across all movie frames, including vesicles visible in the TIRFM zone for at least three frames. Images with mild fading were auto-thresholded in Imaris to ensure consistent tracking.

### Hematopoietic stem and progenitor cell (HSPC) isolation, transplantation, and engraftment

Bone marrow cells were extracted from the femurs of 6- to 8-week-old *Ctns^-/-^* or WT-GFP^+^ transgenic mice. HSPCs were isolated through immunomagnetic separation utilizing an anti-Sca1 antibody linked to magnetic beads (Miltenyi Biotec). The ∼2 × 10^6^ Sca1^+^ HSPCs cells suspended in 100 uL of phosphate-buffered saline (PBS) were then directly injected via tail vein into previous day lethally irradiated (7 Gy; X-Rad 320, PXi) *Ctns^-/-^* mice. In the case of mice receiving WT GFP^+^ HSPCs, engraftment of the transplanted cells was assessed in peripheral blood two months post-transplantation. Blood samples obtained from the tails were treated with red blood cell lysis buffer (eBioscience) and subsequently examined using flow cytometry (BD Accuri C6, BD Biosciences) to ascertain the proportion of GFP^+^ cells. Blood engraftment % is presented in Table S4.

### Immunofluorescence, image acquisition and analysis

Murine kidney tissue sections were collected from euthanized mice, fixed using 10% neutral buffered formalin (NBF), and then embedded in paraffin wax. Standard methods were employed to section the tissue at a thickness of 5 μm. After deparaffinization, the sections were transferred to pre-warmed antigen retrieval solution at 95°C for 30 minutes, followed by cooling at room temperature for 20 minutes. Subsequently, the sections were placed in blocking solution (0.25% Triton X-100 and 3% BSA in tris-buffered saline) and then incubated overnight at 4°C with primary antibodies: rabbit/rat anti-NHE3 (1:50; BiCell Scientific) and rabbit anti-GFP (1:500; Abcam). The next day, appropriate Alexa Fluor-conjugated secondary antibodies (Invitrogen) were added for antigen visualization, along with LTL Rhodamine conjugate (1:100; BiCell Scientific) and DAPI stain. Images were captured using a Keyence BZ-X710 digital microscope. ImagePro Premier software (Media Cybernetics) was utilized for all quantification with thresholding, determining the proportion of expression and colocalization (Pearson’s correlation coefficient) for NHE3 and LTL in z-stacks (multiple images taken per section) or stitched images. Regarding Immunocytochemistry (ICC), the cells underwent the following treatments: 4% paraformaldehyde fixation, 0.5% Triton-X-100 permeabilization, and blocking with 2.5% BSA in PBS, following established protocols thereafter.

NHE3-GFP transduced PTCs were seeded at 70% confluence in a 96-well plate with glass-like polymer bottom black frame (P96-1.5P, Cellvis), then fixed with 4% paraformaldehyde (Electron Microscopy Sciences, 15,710,) for 8 min and blocked with 1% BSA (Rockland, BSA-50) in PBS (Corning, 21–031-CV), in the presence of 0.01% saponin (Calbiochem, 558,255), for 1 h. Samples were labeled with the indicated primary antibodies overnight at 4°C in the presence of 0.01% saponin and 1% BSA. Samples were washed 3 times and subsequently incubated with the appropriate combinations of Alexa Fluor (488 or 594)-conjugated anti-goat, or anti-mouse secondary antibodies (Thermo Fisher Scientific, A-32814, A-21203, respectively). Nuclei were stained with Hoechst 33342 (DAPI; Millipore-Sigma, D9542) and samples were preserved in Fluoromount-G reagent (AnaSpec, AS-83218) and kept at 4°C until analyzed. Samples were analyzed with a Zeiss LSM 880 or Zeiss LSM 980 laser-scanning confocal microscope attached to a Axio Observer Z1 microscope at 21°C, using a 63x oil Plan Apo, 1.4-numerical aperture objective. Images were collected using ZEN-LSM software and processed using ImageJ and Adobe Photoshop. The laser power and gain were maintained throughout the experiments to analyze wild-type and *Ctns^-/-^* cells comparatively. Images were collected and fluorescence intensity and colocalization were quantified using ZEN-LSM software. The following antibodies were used for immunofluorescence in this study: anti-LAMP1 (Santa Cruz Biotechnology, sc-19,992); anti-EEA1 (BD Transduction Laboratories™, 610457); anti-GM130 (BD Transduction Laboratories™, 610822) goat anti-GFP (SICGEN, AB0020-200).

For confocal images, all were acquired using the full dynamic intensity range (1-16383/1-65535) of the specified fluorophores. For quantitative colocalization analysis, regions of interest were drawn around individual cells. Background thresholds were established using secondary antibody controls and further refined using ImageJ’s threshold tool to distinguish specific signal from background. These thresholds were typically set between 500-800 (in the 16-bit dynamic range of 0-65536). The weighted colocalization coefficient (Mander’s overlap coefficient) was calculated as MOC = ∑i(Ri × Gi)/√(∑iRi² × ∑iGi²), where Ri and Gi represent the intensity values of corresponding pixels in the red and green channels.

